# Nutrient-dependent trade-offs between ribosomes and division protein synthesis control bacterial cell size and growth

**DOI:** 10.1101/2020.03.08.982728

**Authors:** Diana Serbanescu, Nikola Ojkic, Shiladitya Banerjee

## Abstract

Cell size control emerges from a regulated balance between the rates of cell growth and division. In bacteria, simple quantitative laws connect cellular growth rate to ribosome abundance. However, it remains poorly understood how translation regulates bacterial cell size and shapes under growth perturbations. Here we develop a whole-cell model for growth dynamics in rod-shaped bacteria that links ribosomal abundance with cell geometry, division control, and the extracellular environment. Our study reveals that cell size maintenance under nutrient perturbations requires a balanced trade-off between ribosomes and division protein synthesis. Deviations from this trade-off relationship are predicted under translational perturbations, leading to distinct modes of cell morphological changes, in agreement with single-cell experimental data on *Escherichia coli*. Furthermore, by calibrating our model with experimental data, we predict how combinations of nutrient-, translational- and shape perturbations can be chosen to optimize bacterial growth fitness and antibiotic resistance.

## INTRODUCTION

Cell size maintenance is essential for regulating cell physiology, function and fitness [1]. Maintaining a characteristic cell size necessitates an intricate balance between cell growth and division rates. How this balance is achieved in different growth conditions remains an outstanding question. It has been known for over six decades that bacteria modulate their size in response to changes in nutrient conditions. Quantifying the cell size and growth rates of *Salmonella enterica* grown in different nutrient media, Schaechter and colleagues discovered the *nutrient growth law* – bacterial cell size increases exponentially with the population growth rate [2]. High-throughput single-cell studies in recent years have confirmed this result for evolutionary divergent *Escherichia coli* and *Bacillus subtilis* [3–6], suggesting common strategies for bacterial cell size control. However, single-cell data show deviations from the nutrient growth law in experiments altering cellular proteomics [4, 7], leaving open the connection between cell size, growth rate and protein synthesis.

At the single-cell level, cell size homeostasis is achieved via the *adder* mechanism, whereby cells add a constant volume between consecutive division events, irrespective of the cell size at birth [5, 8–10]. As a result of this strategy, cells deviating from the average homeostatic size quickly converge to the average size within a few generations [5, 11–14]. This strategy for cell size homeostasis is followed by a wide range of bacterial species including *E. coli, B. subtilis, C. crescentus* and *P. aeruginosa* [5, 11, 13–15], but it does not reveal a molecular-level understanding of the mechanism for cell size control [16].

Two distinct types of regulatory models have been proposed in recent years for the control of bacterial cell size – (1) replication-initiation-centric model: cell size control is set by the time period of chromosome replication and the subsequent cell division (*C* + *D* period) [8, 17, 18], and (2) division-centric model: cell size is regulated by the accumulation of a threshold amount of cell envelope precursors [19] or division proteins (e.g. FtsZ) [20, 21]. In replication-initiation-centric models, cell size at division is determined by the *C* + *D* period, but it remains poorly understood how *C* and *D* periods are modulated by growth perturbations targeting translation or protein expression. Recent studies have challenged the replication-initiation-centric models for cell size control [20, 22–24], suggesting concurrence of replication initiation and division processes [23]. In particular, experiments have demonstrated that replication initiation and cell division are independently controlled in *E. coli* and *B. subtilis* [20], and data support a model that cell division is triggered by the accumulation of a threshold amount of division proteins [13, 25]. However, it remains unknown how the synthesis of division proteins is altered by nutrients or translational perturbations in order to regulate cell size.

A key component in understanding cell size regulation is the interdependence between growth rate and the macromolecular composition of the cell. The nutritional content of the growth medium sets the specific growth rate [2, 26], which in turn regulates the macromolecular composition of cells [27, 28]. For exponentially growing *E*.*coli* cells, RNA and ribosome abundance increases linearly with increasing the specific growth rate [7, 29–33]. This implies an upregulation in translation leading to increased protein production for growth [7, 34, 35] and cell size inflation. While this model is in agreement with experimental observations for cell size increase with increasing nutrient concentrations, it fails to explain cell size changes under translation inhibition [4, 7]. In particular, it remains unclear whether translation inhibition would lead to an increase in cell size such that there is a positive correlation between cell size and ribosome abundance, or a decrease in cell size with growth rate reduction. Both these behaviors are observed in experiments [4]. To explain how translation and nutrient quality regulates cell morphologies, we develop a whole-cell coarse-grained theory that links ribosomes with cell geometry, division control and the extracellular environment.

Our theoretical framework combines a mechanistic model of cell shape and division with an extended ribosomal resource allocation model, allowing us to quantitatively predict cell size changes under nutrient shifts and translational perturbations. We use ribosome abundance as the one of the key regulatory variable as approximately 85% of cellular RNA encodes for rRNA that is folded in ribosomes [36, 37]. We also assume that all the nutrients transported from the extracellular medium into the cell are used in the production of ribosomes and other proteins. This is because over 80% of cell’s energy budget for biomass is spent on rRNA and protein synthesis [38]. Using this framework, we uncover a model for balanced allocation of ribosomal resources towards cell growth and division. We find that a balanced trade-off between the rates of cell growth and division proteins synthesis sets bacterial size under nutrient shifts. As a result, in rich media, cells produce division proteins slower than the rate of cell elongation leading to larger cell sizes.

We then extend our framework to predict cell morphologies under translation inhibition across different nutrient media. Our model predicts three different types of cell morphological response unifying past experimental observations [4, 7, 36]. First, cells deprived of nutrients allocate more ribosomes towards growth which results in an increase in volume. Second, cells grown in rich nutrient media favor resource allocation towards division and thus a decrease in volume is observed under translation inhibition. Under optimal growth conditions cells preserve the balance between growth and division protein synthesis, such that cell size is invariant under translational perturbations. We show that cell size changes are intimately coupled to the regulation of cell surface-to-volume ratio – a fitness metric that controls nutrient and antibiotic influx rates. We therefore investigate the relationship between cell shape, nutrient quality and bacterial growth rate under translational perturbations. We predict that round cells are most resistant to translation-inhibitory antibiotics, and that drug resistance increases with increasing nutrient quality. Thus, induced filamentation could have a negative impact on bacterial growth fitness [39], whereas cell rounding could promote bacterial resistance to ribosome-targeting antibiotics.

## RESULTS

### Cell size control emerges from nutrient-dependent trade-off between rates of cellular growth and division protein synthesis

To understand how bacterial cell size changes with the nutrient specific growth rate, we develop a model for the allocation of ribosomal resources towards cell growth and division protein synthesis. During each cell cycle, cells elongate exponentially in volume (*V*) at a rate *κ*. At steady-state, *κ* depends linearly on the ribosomal mass fraction *r* (≈ RNA/protein ratio), such that

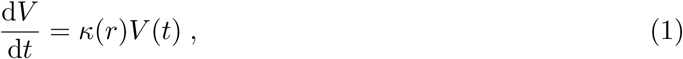

where *κ*(*r*) = *κ*_*t*_(*r* − *r*_min_) [36]. Here, *κ*_*t*_ can be interpreted as the translational capacity of the cell, which correlates with the speed of translational elongation [40], and *r*_min_ is the minimum mass fraction of ribosomes needed for growth (Figure 1C inset). The value for *r*_min_ is obtained from the intercept of *κ* as a function of *r* from experimental data [4, 36].

**Figure 1.**
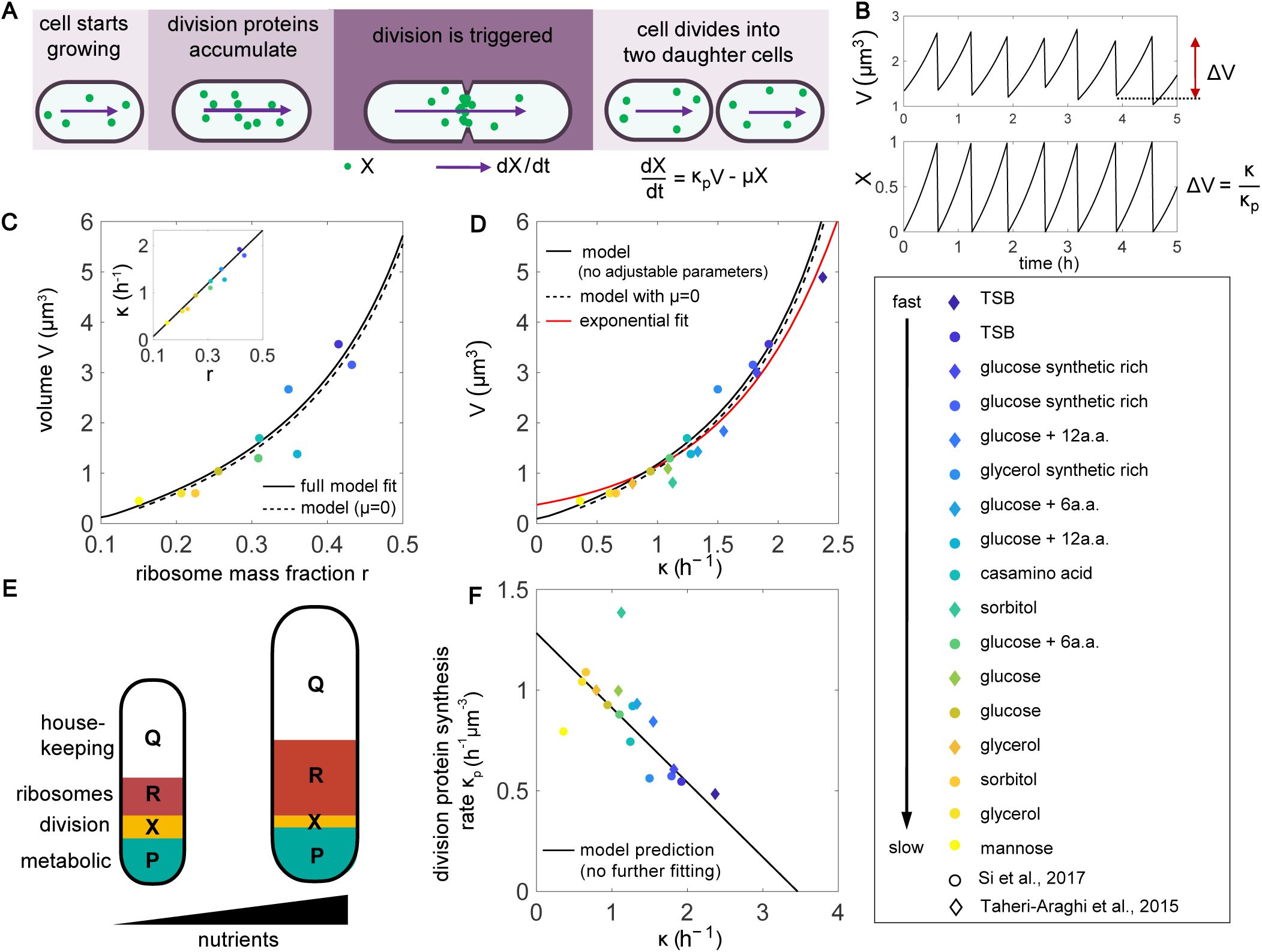
Cell size control under nutrient perturbations. **A**. Schematic of the threshold initiation model. Once a threshold amount of division proteins, *X*, is accumulated, cell division is triggered. Division proteins are synthesized at a rate *κ*_*p*_ per unit volume. **B**. Dynamics of cell volume and fraction of division proteins show that the average added volume between consecutive division events is constant, consistent with the phenomenological adder model [4]. **C**. Fitted model for average cell volume as a function of ribosome mass fraction. Solid line: Full model fit with X protein degradation, Dashed line: Approximate model with no X degradation (*μ* = 0). Inset: fitted linear relationship *κ* = *κ*_*t*_(*r* − *r*_min_) [36]. **D**. Model prediction for the relationship between the average volume and growth rate compared against an exponential fit as predicted by the nutrient growth law [2]. **E**. Schematic representation of the proteome allocation model, showing ribsomal tradeoff between cell growth and division protein synthesis for cells growing in poor and rich nutrient media. **F**. Negative correlation between the rate of division protein synthesis and growth rate, as predicted by the model. Experimental data are obtained from Si *et al*. [4] and Taheri-Araghi *et al*. [5]. In the experimental data *κ*_*p*_ is estimated from the ratio *κ/*⟨*V*⟩. See Table 1 for a complete list of parameter values.

We combine this model for growth with a model for the control of cell division (Figure 1A). The division proteins, *X*, are synthesised at a rate proportional to the cell volume and degraded at a rate *μ*:

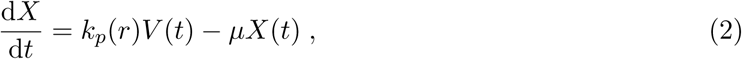

where *k*_*p*_(*r*) is the rate of synthesis of division proteins that is assumed to be a function of the ribosome mass fraction *r*. Cell division is triggered when a threshold copy number of division proteins, *X*_0_, is accumulated at the mid-plane of the cell (Figure 1B). While various proteins could be potential candidates for division initiation [41–44], a recent study identifies FtsZ as the key initiator protein that assembles a ring-like structure in the mid-cell region to trigger septation [20]. We therefore suggest that *X* represents FtsZ copy number, and assume that its turnover rate in the ring-bound state is much faster than its rate of synthesis [45]. As a result, all the newly synthesised FtsZ in the cytoplasm are assumed to be recruited in the ring. We note that the degradation of *X* is consistent with reports of active degradation of FtsZ by ClpXP [20, 46, 47].

**Table 1.**
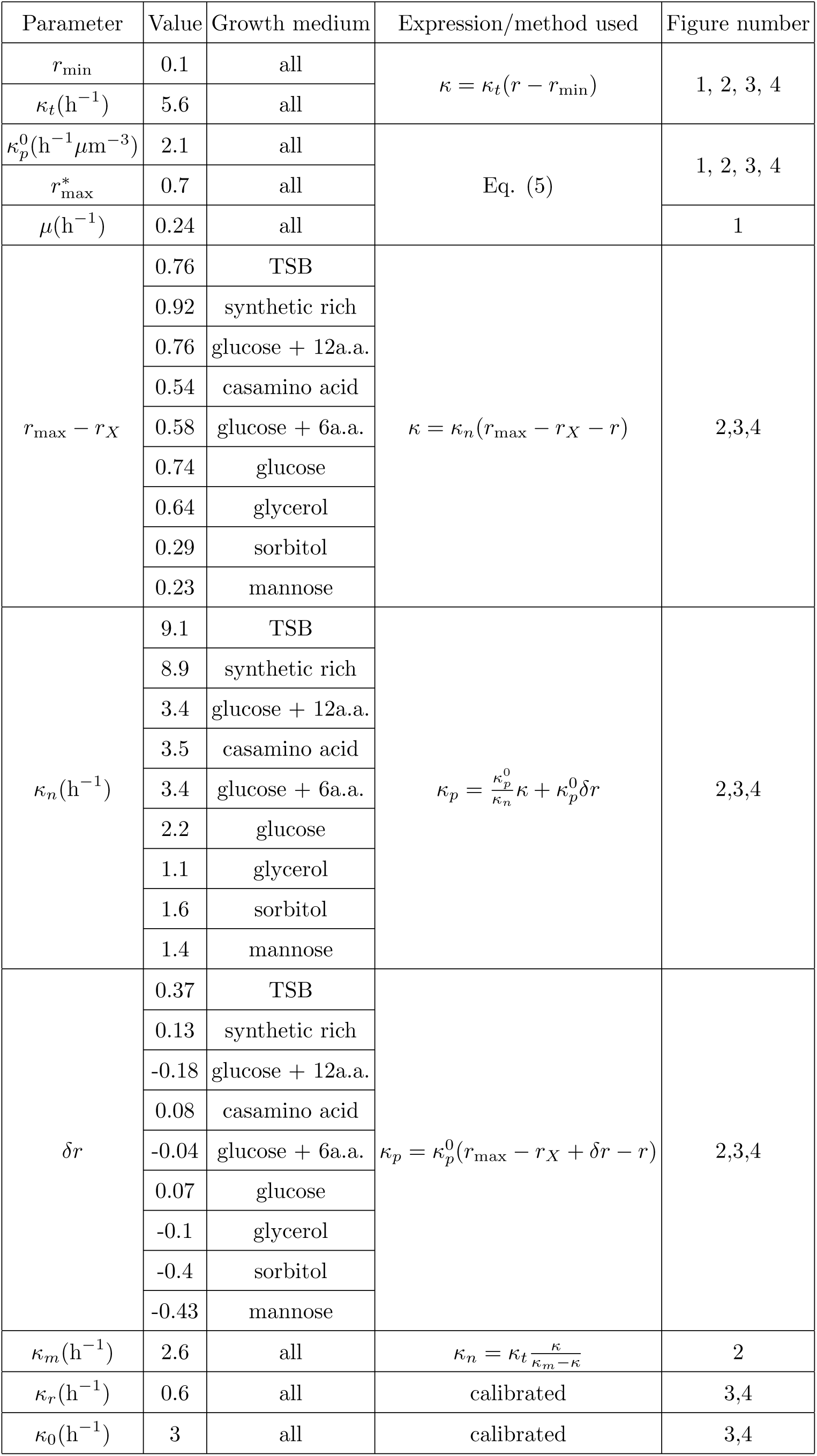
List of parameters used in models of growth perturbations (Figures 1-4).

Solving Eq. (1) and Eq. (2), we obtain: *X*_0_ = (*V*_d_ − *V*_b_2^−*μ*/*κ*^)*k*_*p*_/(*κ* + *μ*), where *V*_b_ and *V*_d_ are the cell volumes at birth and division, respectively. In the limit *k* ≫ *μ*, we get *X*_0_ = Δ*V k*_*p*_*/κ*, where Δ*V* = *V*_d_ − *V*_b_ is the added volume per generation. As *X*_0_, *k*_*p*_ and *κ* are constant for a given growth medium, cells add a constant volume Δ*V* in each growth generation, consistent with the phenomenological adder model. Conversely, in the limit *k* ≪ *μ, V*_d_ ≈ *X*_0_*μ/k*_*p*_, consistent with data that *E. coli* deviates from an adder in slow growing media [8, 20]. Furthermore, for symmetrically dividing bacterium, the average newborn cell volume, ⟨*V*_b_⟩, asymptotes to Δ*V* [11]. Therefore, average cell volume ⟨*V*⟩ in a given growth medium is given by:

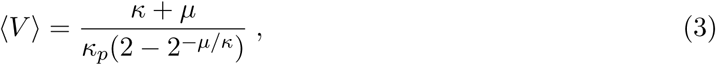

where *κ*_*p*_ = *k*_*p*_/(2*X*_0_ ln 2), is the normalized rate of division protein synthesis. Thus, cell volume can be modulated by perturbations in translation, as both *κ* and *κ*_*p*_ are functions of the ribosomal mass fraction. A key proposition of our model is that there is a tradeoff between ribosomes allocated for synthesizing growth and division proteins such that:

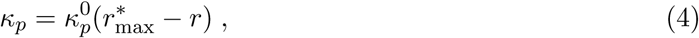

where we interpret 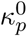 as the rate of production of FtsZ per ribosomes, and 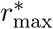 is the ribosome mass fraction when growth rate is maximum. We note that the parameter 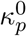 can be perturbed by translation, whereas 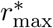 is regulated by both translational and nutrient capacities of the cell, as derived in the following section (see STAR Methods for further details). By combining the expressions for growth rate and division proteins synthesis rate, we find:

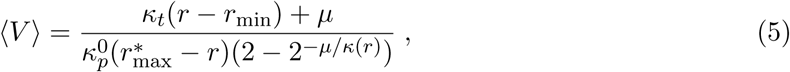

such that average cell size increases with increasing ribosome abundance. We fit the expression in Eq. (5) to experimental data [4] in order to determine the parameters 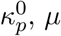, and 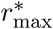 (Figure 1C, solid line). Importantly, we find that *μ* = 0.24 h^−1^, allowing us to approximate the average volume as the ratio of growth rate to the rate of division protein synthesis: 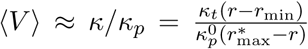 (Figure 1C, dashed line). Thus *κ*_*p*_ can be indirectly measured from *κ/*⟨*V*⟩ data across different growth conditions. Direct measurement of *κ*_*p*_ would necessitate measuring the rate of change in FtsZ fluorescence intensity per unit cell volume during cell division cycles.

We can then express the average cell volume as a function of nutrient specific growth rate, recapitulating Schaecter et al.’s *nutrient growth law* [2] that cell size increases monotonically with increasing growth rate:

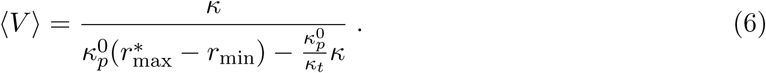

With no further fitting, we directly compare the prediction in Eq. (6) with experimental data for *E. coli* cell volume under nutrient perturbations (Figure 1D). The result in Eq. (6) deviates from the phenomenological model of exponential dependence between cell size and growth rate, and predicts a maximum growth rate, 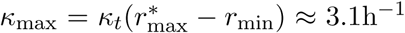, when all ribosomal resources are allocated towards growth. Our model captures the departure from an exponential relationship between cell size and growth rate for *κ* < 0.7 h^−1^, as recently reported by Zheng et al [48] (Figure S1A). We also find that a linear relationship between cell size and growth rate does not accurately capture the cell size data for the range of growth rates studied in this work.

### Mechanistic origin of ribosomal tradeoff between growth and division

To understand the mechanistic origin of the ribosomal tradeoff between growth and division protein synthesis (Eq. 4), we develop a model for allocation of ribosomal resources, extending the framework of Scott et al [36]. The total protein content of the cell can be decomposed into four classes (Figure 1E): ribosome-affiliated proteins (R, mass fraction *ϕ*_*R*_), house-keeping proteins not affected by translation (Q, mass fraction *ϕ*_*Q*_), division proteins (X, mass fraction *ϕ*_*X*_), and the rest non-ribosomal proteins that constitutes the metabolic sector (P, mass fraction *ϕ*_*P*_). The mass fractions are constrained by the equation: 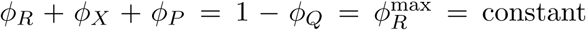. For different combinations of the nutritional and translation capacities of the cell, efficient resource allocation requires that the abundance of P- and R-class proteins be adjusted so that the rate of nutrient influx by P matches the rate of protein synthesis achievable by R: 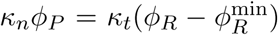, where *κ*_*n*_ is the nutritional capacity of the cell. This results in the following relation between the mass fractions of ribosomes and division proteins:

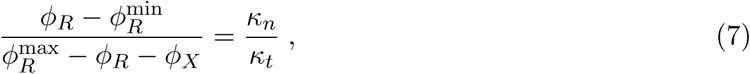

predicting a negative correlation between *ϕ*_*X*_ and *ϕ*_*R*_ under nutrient or translational perturbations (Figure S1C). Using a dynamic proteome sector model (STAR Methods) we can derive that the rate of production of division proteins, *κ*_*p*_, is proportional to *ϕ*_*X*_ during steady-state growth. Using *r* = *ϕ*_*R*_/*ρ*, where *ρ* is a constant conversion factor, we derive the negative correlation between *κ*_*p*_ and *r*, as assumed in Eq. (4),

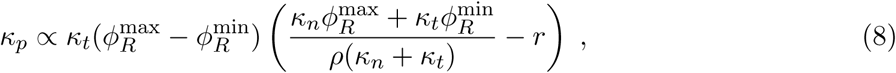

where we identify 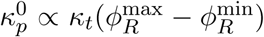 and 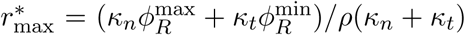 (see details in STAR Methods). Thus the tradeoff between ribosomes, growth and division protein synthesis naturally emerges in an extended proteome allocation model. Growth rate *κ* decreases with increased allocation of resources towards division proteins *ϕ*_*X*_ (Figures 1E-F and S1D):

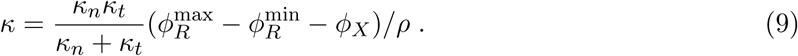

With all the model parameters inferred from experimental data (STAR Methods, Table 1), we can plot the dependency of *κ*_*p*_ on *κ* (Figure 1F), showing the negative correlation between the division protein synthesis rate and the volumetric growth rate, and directly predict the dependency of cell volume on growth rate (Figure S1B). Cells growing in poor nutrient medium allocate a smaller fraction of ribosomes towards growth, resulting in smaller size on average. However, cells growing in rich nutrients inflate their size by allocating a larger fraction of ribosomes towards growth (Figure 1E).

### Translation inhibition breaks balanced allocation of ribosomal resources

In a given nutrient medium, *κ/κ*_*p*_ is maintained at a constant value, indicating a balance between growth and division protein synthesis. If *κ/κ*_*p*_ remains invariant under translation inhibition, we expect cell size to remain unchanged, as previously suggested by Basan *et al*. [7]. However, experimental data [4] show that cell size could either increase, decrease or remain unchanged when *E. coli* cells are subjected to varying concentrations of Chloramphenicol – a ribosome-targeting antibiotic. We therefore hypothesize that translation inhibition breaks balanced allocation of ribosomal resources towards growth and division proteins, by differentially reducing the rates *κ* and *κ*_*p*_.

Under translation inhibition, bacteria produce more ribosomes to compensate for the inactive ribosomes that are bound by antibiotics [36]. By measuring bacterial growth rates (*κ*) and ribosome mass fractions (*r*) for increasing concentrations of Chloramphenicol, Scott *et al*. [36] found that *κ* linearly decreases with *r*. In the presence of a division protein sector, the relationship between *κ* and *r* is given by:

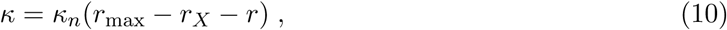

where *κ*_*n*_ is the nutritional capacity that depends on nutrient quality, 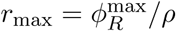 is the maximum ribosome fraction that cells can produce under translation inhibition, and *r*_*X*_ = *ϕ*_*X*_/*ρ* is the RNA/protein ratio devoted to synthesizing *X* proteins. By combining Eq. (10) with Eq. (4), we obtain:

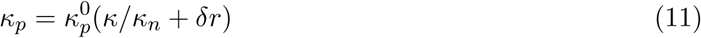

where 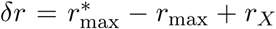 can be interpreted as the excess ribosomal mass fraction allocated to division protein synthesis under translation inhibition. By combining Eqs. (4), (7), and (10) we obtain a theoretical expression for the excess ribosomal mass fraction as a function of the growth rate: *δr* = *κ*_*n*_Δ*r/*(*κ*_*n*_ + *κ*_*t*_) − *κ*(*κ*_*n*_ + *κ*_*t*_)/(*κ*_*n*_*κ*_*t*_), where Δ*r* = *r*_max_ − *r*_min_. Since *κ*_*n*_ increases with *κ* (Figure 2C-inset), we predict that *δr* increases monotonically with nutrient specific growth rate (Figure 2D, solid line).

**Figure 2.**
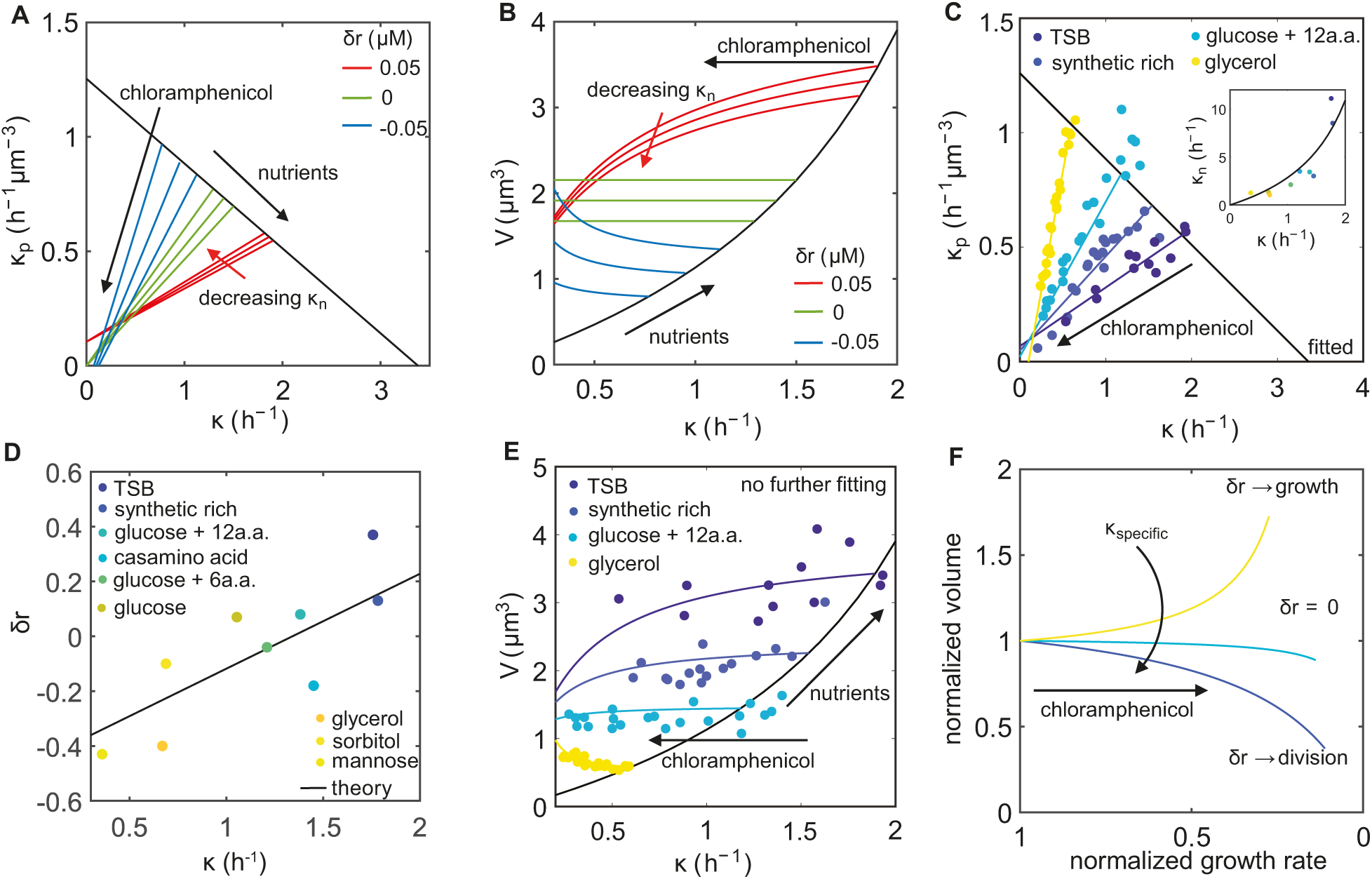
Cell size control under translation inhibition. **A**. Model prediction for the dependence of division protein synthesis rate, *κ*_*p*_, on growth rate, *κ*, under translation inhibition for three different values of *δr*. Decreasing *κ*_*n*_ corresponds to increasing the nutrient quality of the growth medium. **B**. Model predictions for cell volume vs growth rate under translation inhibition, capturing three distinct trends in cell size changes depending on the value of *δr*. **C**. Model fit to experimental data for *κ*_*p*_ as a function of *κ* in three different nutrient conditions under translation inhibition. Inset: Dependence of nutritional capacity *κ*_*n*_ on growth rate. Solid line is a fit of the form: *κ*_*n*_ = *κ*_*t*_*κ/*(*κ*_*m*_ − *κ*), with the fitting parameter *κ*_*m*_ = 2.6 h^−1^. **D**. Dependence of *δr* on nutrient specific growth rate. Solid line shows the theoretical prediction for the dependence of *δr* on nutrient specific growth rate, 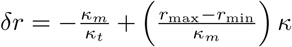. **E**. Cell volume changes with growth rate under translation inhibition. **F**. Schematic illustrating three distinct morphological response to Chloramphenicol, depending on the quality of nutrients. In nutrient-rich media, cells allocate more ribosomes to division (dark blue line) thus increasing the surface-to-volume ratio to promote nutrient influx, while in nutrient-poor media they allocate the more ribosomes towards growth, inflating the cell size (yellow line) and in turn decreasing the surface-to volume ratio to reduce the antibiotic influx. See Table 1 for a complete list of parameter values.

Unlike nutrient perturbations, we find that *κ*_*p*_ and *κ* are positively correlated under translation inhibition (Figure 2A), such that they both decrease with increasing antibiotic concentration. Eq. (11) can be combined with Eq. (3) to determine how cell volume changes as a function of growth rate under translation inhibition:

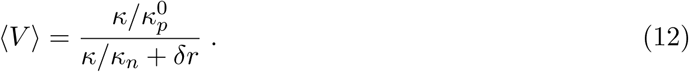

Interestingly, the above expression predicts three distinct behaviors (Figure 2B):

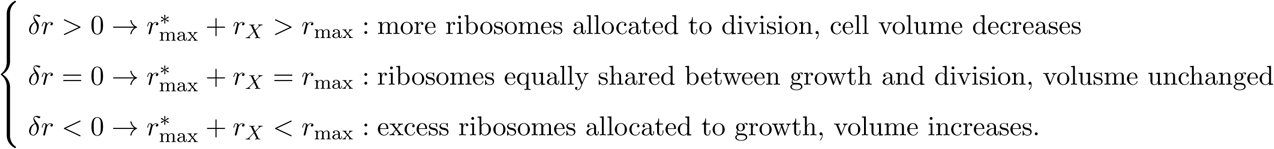

We determine the parameters *δr* and *κ*_*n*_ for each growth medium, by fitting Eq. (11) to the experimental data for *κ*_*p*_ vs cell growth rate *κ* under Chloramphenicol perturbations [4] (Figure 2C). We find that *δr* < 0 in poor media, *δr* > 0 in rich growth media, whereas *δr* ≈ 0 for cells growing with medium growth rates (Figure 2D). These data are consistent with our theoretical result that *δr* increases linearly with the nutrient specific growth rate (Figure 2D-solid line). We interpret the above result as cells allocating excess ribosomes for growth in poor media, whereas in rich media cells tend to allocate more ribosomal resources for division protein synthesis. With no further adjustable parameters, our theory predicts the cell volume curves for each growth conditions, which are in excellent quantitatively agreement with the trend in the experimental data (Figure 2E).

### Cells actively regulate shapes to adapt to translational perturbations

Under translation inhibition, decrease in cell volume in rich media is indicative of a higher surface-to-volume ratio that may increase the influx of nutrients and antibiotics. Conversely, in poor media, increase in cell volume may be indicative of a lower surface-to-volume ratio that in turn would reduce antibiotic and nutrient influx (Figure 2F). Therefore, surface-to-volume ratio of a cell may play a crucial role in controlling cellular adaptive response to growth perturbations, by modulating the relative contributions of nutrient and antibiotic influx rates. To test the role of surface-to-volume ratio on bacterial growth, we construct a model coupling cell growth and geometry to nutrient and antibiotic transport.

#### Nutrient dynamics

The dynamics of nutrient concentration inside the cell, [*n*], is given by:

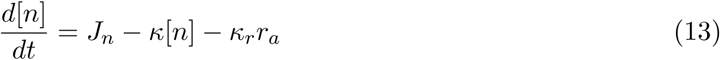

where *J*_*n*_ = [*n*_ext_]*P*_in_*A/V* is the nutrient influx, [*n*_ext_] is the nutrient concentration in the extra-cellular medium, *P*_in_ is the cell envelope permeability and *κ*_*r*_ is the rate at which ribosomes are produced from the nutrients. The model for nutrient transport across the cell membrane is consistent with the one proposed in [49] if we assume that the number of metabolic proteins (transporters) scales with the surface area of the cell. The interplay between nutrients and ribosome synthesis is schematically represented in Figure 3A. The intracellular concentration of nutrients determines the specific growth rate as: *κ*_specific_ = *κ*_0_[*n*]/([*n*] + *n*^∗^) [50], where *κ*_0_ is the maximum growth rate characteristic of the medium, and *n*^∗^ is the value of [*n*] when *κ*_specific_*/κ*_0_ = 0.5. When the nutrients inside the cell reach saturation, i.e. d[*n*]/d*t* = 0, *κ* = *κ*_specific_.

**Figure 3.**
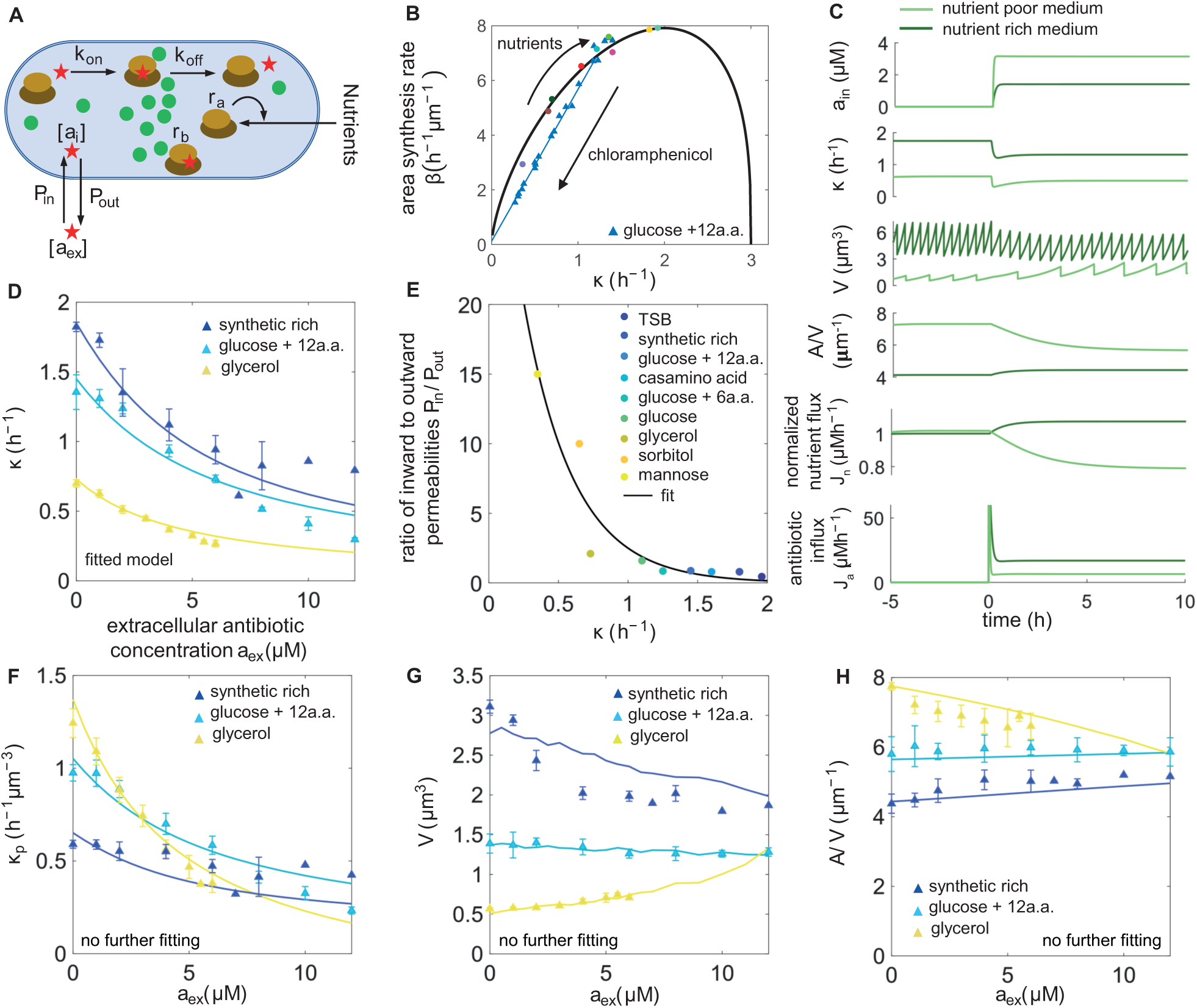
Cell shape changes under translation inhibition. **A**. Schematic illustrating nutrient and antibiotic transport across cell surface and antibiotic interactions inside the cell. **B**. Model predictions for the surface area synthesis rate (*β*) as a function of the growth rate (*κ*) for varying nutrient conditions, and its inhibition under Chloramphenicol perturbations. *β* is calculated using *β* = *κA/V*. **C**. Single-cell simulations of growth in response to a step pulse of Chloramphenicol applied at *t* = 0 h in the extracellular medium. Top to bottom: Dynamics of intracellular antibiotic concentration, growth rate, cell volume, surface-to-volume ratio, normalized nutrient and antibiotic flux. Both nutrient and antibiotic fluxes are higher in rich media due to the increase in surface-to-volume ratio. **D**. Model growth inhibition curves fitted to experimental data [4], in order to deduce the ratio of inward to outward cell-surface permeability. Data are represented as mean ± SEM. **E**. *P*_in_*/P*_out_ increases with decreasing growth rate (quality of nutrients). We used an exponential fit to calibrate *P*_in_*/P*_out_ for different nutrient conditions. **F-H**. Simulation results for the dependence of *κ*_*p*_ (F), average cell volume *V* (G), and average cell surface-to-volume ratio *A/V* (H) on Chloramphenicol concentration, plotted against experimental data [4] at three different nutrient conditions. Data are represented as mean ± SEM. See Tables 1 and 2 for a complete list of parameter values.

#### Antibiotic dynamics

The action of ribosome-targeting antibiotics is illustrated using the diagram in Figure 3A, which consists of two key components: the flux of antibiotics *J*_*a*_ entering the cell, and the binding of antibiotics to the active pool of ribosomes, *r*_*a*_. The dynamics are described by the following set of equations, extending the model of Elf *et al*. [51] and Greulich *et al*. [52],

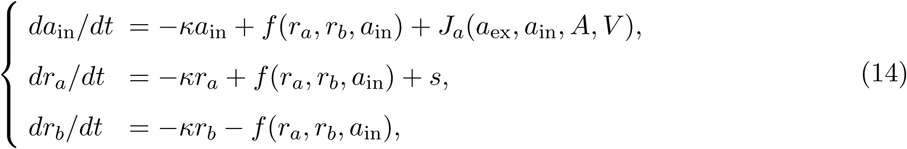

where *a*_ex_ is the extracellular antibiotic concentration, *a*_in_ is the intracellular concentration of the antibiotic, *r*_*a*_ is the concentration of the active pool of ribosomes in the cell, *r*_*b*_ is concentration of the pool of ribosomes bound by the antibiotics, and *s* is the rate of synthesis of ribosomes. Unlike previous models [51, 52], here we account for the dependence of *J*_*a*_ on cell shape as:

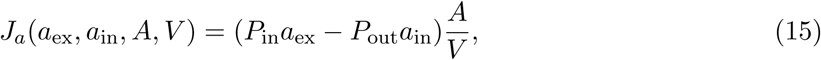

where *P*_in_ and *P*_out_ are the cell envelope permeabilities in the inward and outward directions, respectively. The ribosome-antibiotic interactions are defined by: *f* (*r*_*a*_, *r*_*b*_, *a*_in_) = −*k*_on_*a*_in_(*r*_*a*_ −*r*_min_)+ *k*_off_*r*_*b*_, where *k*_on_ is the rate of binding of antibiotics to ribosomes, and *k*_off_ is the rate of unbinding. These rate constants for Chloramphenicol are known from literature [51, 52]. Furthermore, cells produce more ribosomes to compensate for the inactive ribosomes bound by antibiotics [36]. This is captured by the source term: *s* = *κ*[*r*_max_ − *κ*Δ*r*(1*/κ*_specific_ − 1*/κ*_*t*_Δ*r*)], where Δ*r* = *r*_max_ − *r*_min_. *Cell shape dynamics*. Having described the dynamics of cell volume (Eq. 1), division control (Eq. 2), nutrient and antibiotic transport (Eq. 13-14), we need to additionally account for cell surface area synthesis to predict cell shape changes. We assume that rate of synthesis cell surface area is proportional to cell volume [19]:

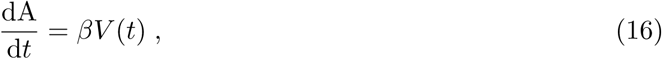

where *β* is the rate of surface area production, which depends on cell shape, growth rate and division protein synthesis rate. Solving Eq. (1) and Eq. (16), one obtains *A/V* = *β/κ*, at steadystate [19]. In recent work [21] we found that *E. coli* cells obey the relation: *A* = *νV* ^2/3^, under nutrient and translational perturbations, where *ν* is a geometric factor related to the cell aspect ratio *η* as: 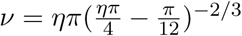. Therefore, surface area production rate varies non-monotonically with growth rate as: *β* = *νκ*(*κ/κ*_*p*_)^−1/3^ (Figure 3B).

**Table 2.** List of parameters used in antibiotic simulations (Figures 3-4).

| Parameter | Value | Growth medium | Expression/method used | Figure number |
| --- | --- | --- | --- | --- |
| $[n_{\text{ext}}]/n^*$ | 0.2 | TSB | Fitting $\kappa = \kappa_0[n]/([n] + n^*)$ when $\frac{d[n]}{dt} = 0$ | 3,4 |
|  | 0.08 | synthetic rich |  |  |
|  | 0.03 | glucose + 12a.a. |  |  |
|  | 0.05 | casamino acid |  |  |
|  | 0.018 | glucose + 6a.a. |  |  |
|  | 0.008 | glucose |  |  |
|  | 0.002 | glycerol |  |  |
| $P_{\text{out}}(\text{h}^{-1}\mu\text{m}^{-1})$ | 20 | all | $15 - 30 \text{ h}^{-1}$ [52] | 3,4 |
| $P_{\text{in}}/P_{\text{out}}$ | 0.45 | TSB | Fitting growth inhibition curves | 3,4 |
|  | 0.80 | synthetic rich |  |  |
|  | 0.88 | glucose + 12a.a. |  |  |
|  | 0.80 | casamino acid |  |  |
|  | 0.85 | glucose + 6a.a. |  |  |
|  | 1.6 | glucose |  |  |
|  | 2.1 | glycerol |  |  |
|  | 10 | sorbitol |  |  |
|  | 15 | mannose |  |  |
| $k_{\text{on}}(\mu\text{M}^{-1}\text{h}^{-1})$ | 10 | all | $1.08 - 13 \mu\text{M}^{-1}\text{h}^{-1}$ [52] | 3,4 |
| $k_{\text{D}}(\mu\text{M})$ | 2.5 | all | $0.5 - 5 \mu\text{M}$ [52] | 3,4 |
| $k_{\text{off}}(\text{h}^{-1}) = k_{\text{D}}k_{\text{on}}$ | 25 | all | calculated | 3,4 |

Taken together, our model accounts for the key functions of ribosomes in controlling cell growth rate (*κ*), rate of production of division proteins (*κ*_*p*_), and the rate of surface area synthesis (*β*). Under translation inhibition, both *κ* and *κ*_*p*_ decreases as shown in Figures 2A and C. Surface area production rate is also impacted by translation inhibition, as shown in Figure 3B, albeit in a different manner from the growth rate. Differential reduction of *κ* and *β* under translation inhibition is indicative of changes in steady-state cell surface-to-volume ratio (∝ *β/κ*). To test this quantitatively, we simulated the coupled equations (STAR Methods) for single-cell growth (Eq. 1, 2 and 16), nutrient (Eq. 13) and ribosome-antibiotic dynamics (Eq. 14) under stresses induced by ribosome-targeting antibiotics (Figure 3C). In response to a step pulse of antibiotic in a rich nutrient medium at *t* = 0 h, the concentration of antibiotic inside the cell and the influx increase rapidly. This in turn reduce cell elongation rate as a result of antibiotic binding to ribosomes, and leads to longer interdivision times, a decreased (increased) average birth volume and a concomitant increase (decrease) in surface-to-volume ratio for cells growing in rich (poor) nutrients. These results confirm our hypotheses that in poor media, cells reduce their surface-to-volume ratio to inhibit antibiotic influx, while in rich media cells increase their surface-to-volume ratio to import more nutrients.

While all the model parameters can be calibrated from available experimental data (Table 1-2, STAR Methods), the relative magnitude of the permeabilities, *P*_in_*/P*_out_ remains undetermined. To this end, we fit our model to the experimental growth-inhibition curves [4] in differents nutrient conditions (Figure 3D), treating *P*_in_*/P*_out_ as a fitting parameter. Interestingly, we find that *P*_in_*/P*_out_ is nutrient-dependent and decreases with increasing specific growth rate (Figure 3E). As a result, *J*_*n*_*/J*_*a*_ is an increasing function of growth rate, such that nutrient influx dominates over antibiotic influx in nutrient rich media. Incorporating nutrient-dependent regulation of membrane permeability, our model predictions capture the experimental data for the decrease in division protein synthesis rate under Chloramphenicol inhibition (Figure 3F) and the changes in cell volume (Figure 3G). Consistent with our hypothesis and experimental data, we find that cell surface-to-volume increases in rich nutrient media (synthetic rich in experiments) (Figure 3H). Conversely, in poor nutrient medium (glycerol in experiments), cell surface-to-volume reduces with increasing drug dosage, suggesting that cells are countering the influx of antibiotics if sufficient nutrients are not available.

Nutrient-dependent regulation membrane permeability to antibiotics (Figure 3E) can be a result of different metabolic pathways. It has been observed that *E*.*coli* cells have different metabolic pathways for nutrients depending on the growth conditions [53]. Furthermore, if the cells are subjected to a nutrient downshift, the proteome reallocates such that a larger fraction of proteins is allocated to the sector responsible for carbon catabolism which in turn reduces the available proteome fraction for other sectors [54, 55]. The transition from one metabolic mechanism to another can be justified using a proteome allocation model as suggested by Basan *et al*. [55] and Mori *et al*. [54], or by increasing the glucose uptake rates. The drop in cell envelope permeability that we observe around *κ* = 0.6 h^−1^ (Figure 3E) matches the maximum growth rate that *E*.*coli* cells can achieve while staying below the critical limit on energy dissipation [56].

### Cell surface area production promotes bacterial growth inhibition by ribosome-targeting antibitiocs

Our theory predicts that bacterial growth response to translation-inhibitory antibiotics is governed by nutrient-dependent cell shape changes (Figures 2 and 3). To systematically study how bacterial growth inhibition depends on cell shape and nutrient quality, we simultaneously perturbed cell shape and ribosomal translation in varying growth media using our computational model. These simulations can be realised experimentally by simultaneously applying two antibiotics - one that changes cell shapes (e.g. by targeting the cell wall), while the other affects the translational machinery by inhibiting ribosomal activity. The resultant effect can be suppressive, antagonistic, or synergistic depending on what the combined effect of the two drugs is with respect to the individual effect of each [57, 58].

In simulations we simultaneously applied a surface area *modifier* and chloramphenicol to a cell growing at steady-state. To achieve rounder cells, the *modifier* is a surface area synthesis inhibitor that decreases the surface production rate *β*, by decreasing the cell’s geometric factor *ν* (= *A/V* ^2/3^), which in turn reduces cellular aspect ratio. By contrast, long filamentous cells are obtained when a surface area promoter is added (increasing *ν*), leading to higher aspect ratio cells.

We investigated the response of growth rate to increasing Chloramphenicol concentrations for cells with varying aspect ratios – ranging from *η* = 1 for coccoidal cells to *η* = 10 for filamentous cells (Figure 4A). The response of *κ* to the concentration of the applied antibiotic can be characterized by a Hill function of the form [59] (Figure 4A):

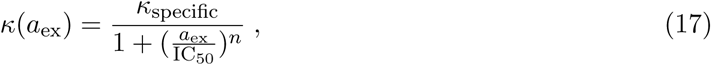

where IC_50_ is the half-inhibitory concentration of the antibiotic, and the Hill coefficient *n* quantifies the dose-sensitivity of the growth rate to relative changes in drug concentration. We take IC_50_ as a measure of drug *resistance* [59].

**Figure 4.**
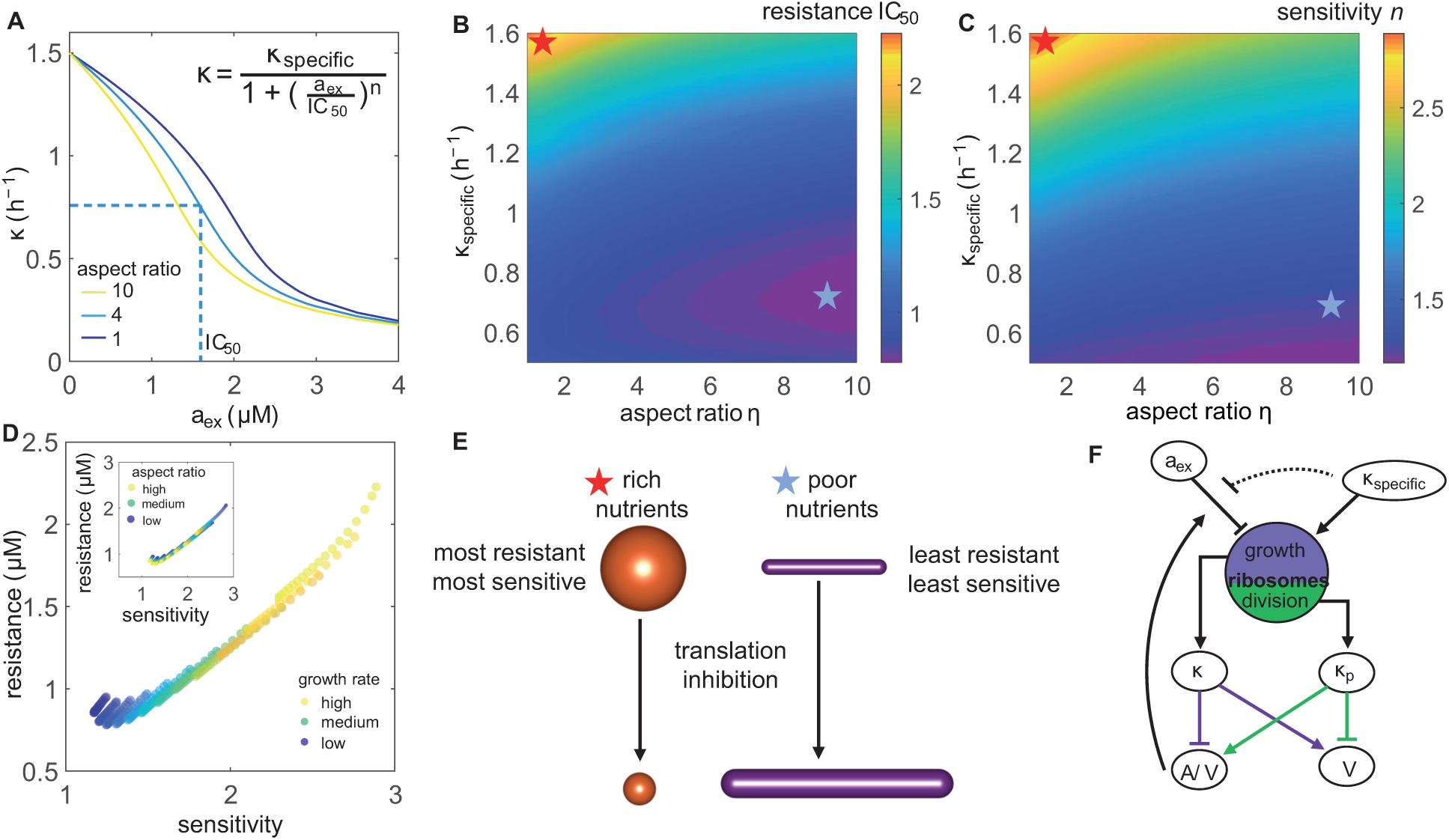
Cell shape and nutrient quality control bacterial resistance to ribosome-targeting antibiotics. **A**. Growth inhibition curves for three different values of cell aspect ratio in a nutrient-rich medium. Dashed line corresponds to IC_50_ on the x-axis, the concentration of antibiotic when the growth rate reduces by half. **B**. Heatmap of IC_50_, a metric for drug resistance, showing the effects of changing aspect ratio and nutrient quality. Red asterix: Maximally resistant, blue asterix: least resistant. **C**. Heatmap of drug dose-sensitivity (*n*), showing the effects of changing cellular aspect ratio and nutrient quality of the growth medium. **D**. Correlation between drug resistance and dose-sensitivity under changes in nutrient quality (*κ*_specific_). Inset: Correlation between drug resistance and dose-sensitivity under changes in aspect-ratio. **E**. Schematic illustrating fitness value for cell shapes and morphological changes that accompany bacterial response to translation inhibition in nutrient-rich and nutrient-poor growth media. **F**. Schematic representation of the feedback pathways that connect ribosomal translation to bacterial cell shape, growth, nutrient and antibiotic transport. See Tables 1 and 2 for a complete list of parameter values.

For a range of aspect ratios and nutrient conditions, we fitted the growth inhibition curves to the Hill function in Eq. (17), and obtained the values for IC_50_ (Figure 4B) and the dose-sensitivity *n* (Figure 4C). Our model predicts that IC_50_ (resistance) increases with decreasing aspect ratio in rich-nutrient medium, while being less sensitive to changes in cell aspect ratio in poor-nutrient medium (Figure 4B). Dose-sensitivity to changes in drug concentration increases with decreasing aspect ratio and increasing nutrient quality (Figure 4C), such that dose-sensitivity is positively correlated with drug resistance (Figure 4D). These results indicate that cellular response to translation inhibitory antibiotics is sensitive to both the nutrient quality as well as cell shape. We find that round coccoidal cells are most drug-resistant, while filamentous cells are least resistant (Figure 4E). Furthermore, depending on nutrient-quality, cellular morphological response to translation inhibitory drugs is different. While cells increase their surface-to-volume ratio to import more nutrients in nutrient-poor medium, cells prefer to reduce their surface-to-volume ratio in rich-nutrient medium to inhibit antibiotic influx (Figures 2-3). These findings predict that bacterial growth inhibition can be maximized by simultaneously inhibiting ribosomal translation and promoting surface area production in nutrient-poor media.

## DISCUSSION

We develop a whole-cell coarse-grained model for bacterial growth dynamics that connects intra-cellular control of translation, with cell shape, division control and extracellular environment. This provides a promising theoretical framework that quantitatively captures available experimental data for bacterial cell size and shape dynamics under nutrient and translational perturbations. Our study reveals that during nutrient shifts, the ribosomal resources are optimally allocated to maintain a balanced trade-off between the rates of cell growth and division protein (FtsZ) synthesis. In rich nutrient media, more ribosomes are used for growth than division protein synthesis, leading to cell size inflation with increasing nutrient quality. Conversely in nutrient-poor media, cells allocate more ribosomal resources for division protein synthesis than growth, leading to a reduction in average cell size. This principle underlies the molecular basis for the celebrated *nutrient growth law* [2, 4], and can be interpreted as an optimization principle for cellular economy. Based on this principle, the resources allocated to a particular proteomic sector are inversely proportional to the efficiency of that sector [49]. In nutrient-rich media, cells invest more ribosomal resources to growth in order to compensate for a lower translational capacity. The latter can arise from an increased dilution rate of ribosomes under fast growth conditions, lowering the efficiency of protein synthesis. In nutrient-poor media, cells have a lower nutritional capacity that they compensate by allocating more resources to metabolism and division protein synthesis.

To explain the mechanistic origin of the ribosomal tradeoff between growth and division protein synthesis, we propose a proteome allocation theory, extending the sector model introduced by Scott et al [36]. In particular, we introduce a division protein sector X in the proteome, and derive a constitutive relation that the mass fraction of X, *ϕ*_*X*_, is a linearly decreasing function of the mass fraction of ribosomes, *ϕ*_*R*_. Existing experimental data support our model, and also falsify other possible models with X in the R-sector or the Q-sector. In particular, if X is the R or Q sector we would expect cell size to always decrease under translation inhibition, a result that is inconsistent with experimental data [4, 7]. A recent study by Bertaux et al [60] also incorporated division protein sector in a proteome model [49]. In contrast to our theory, the authors assumed a phenomenological form for the dependence of *ϕ*_*X*_ on *ϕ*_*P*_ and 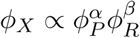, where the exponents *α* and *β* are deduced by fitting experimental data. Bertaux et al’s findings are consistent with our results for cell size control under nutrient perturbations.

Our model for division control is chromosome agnostic, and is thus inadequate for capturing the single-cell correlation patterns related to DNA replication initiation and segregation periods [23, 24]. However, we find that the rate of production of division proteins, *κ*_*p*_, is proportional (*C* +*D*)^−1^ (Figure S2A-B), where *C* is the duration from initiation to termination of one round of DNA replication, and *D* is the time period from replication termination to cell division. This relationship between adder protein synthesis and chromosome dynamics emerges from combining the principle of balanced biosynthesis *κ*_*p*_ ∝ *κ* [20] with the relation *C* + *D* ∝ *κ*^−1^ [61]. The proportionality *κ*_*p*_ ∝ (*C* + *D*)^−1^ also emerges from the recent model suggested by Zheng et al that cell size is linearly proportional to *κ*(*C* + *D*) [48]. Using *V* ∝ *κ*(*C* + *D*) in conjunction with our theory *V* ∝ *κ/κ*_*p*_ also reveals *κ*_*p*_ ∝ (*C* + *D*)^−1^, consistent with experimental data [20]. However, we note that a linear relationship between cell size and *κ*(*C* + *D*) does not accurately describe all available experimental data [4, 7, 62] for cell size (Figure S2C).

Comparing our theory to experimental data, we uncover several feedback pathways between cell shape, growth rate, protein synthesis and extracellular transport that were previously unknown (Figure 4F). In particular, we predict that under translation inhibition, cells break the balanced trade-off between ribosomes and division protein synthesis, leading to cell size inflation, reduction or size invariance, in a nutrient-dependent manner. The model proposed by Bertaux et al [60] predicts that cell size always increases under translation inhibition, unless the cells followed a sizer-like law for cell size homeostasis. Our model predictions are in quantitative agreement with experimental data on *E. coli* cells subjected to Chloramphenicol perturbations across various nutrient conditions [4]. If cells are grown in nutrient-rich media, the excess ribosomes produced under translation inhibition are allocated towards division, leading to smaller cell sizes and higher surface-to-volume ratios. This is in agreement with Chloramphenicol treated *E*.*coli* cells grown in synthetic rich medium. Conversely, in nutrient-poor media cells allocate excess ribosomes towards growth, leading to cell size inflation and lower surface-to-volume ratios, in agreement with *E*.*coli* cell data in glycerol medium.

Our results suggest that cells shape changes in response to translation-inhibitory antibiotics may confer certain fitness advantages under stress. In nutrient-rich media it is more favorable for cells to reduce their surface-to-volume in order to minimize antibiotic influx. Whereas in nutrient-poor media, cells adapt to import more nutrients by increasing their surface-to-volume ratios. To quantitatively test the role of cell shape and nutrient quality on bacterial growth inhibition under antibiotic stress, we simulated bacterial growth under simultaneous perturbation of surface area production and translation inhibition in varying nutrient media. From growth-inhibition curves we measured bacterial response to antibiotics by quantifying resistance (half-inhibitory concentration of the drug) and dose-sensitivity to increasing concentration of the drug. Our study reveals that that round-shaped cells are fitter and more drug-resistant than higher aspect ratio filamentous cells, and that dose-sensitivity increases with increasing nutrient quality. These results can be tested experimentally by measuring bacterial growth rates in response to simultaneous application of cell-wall targeting and ribosome-targeting antibiotics, in different nutrient concentrations. Interestingly, we predict that bacterial growth-inhibition can be maximized by simultaneously inhibiting ribosomal translation and promoting surface area production in nutrient-poor media.

## METHODS

### Cell growth simulations

To investigate the dynamic response of cell shape and growth to applied antibiotic and nutrient shifts, we simulated single-cell growth over multiple generations. We first initiated cells at random stages in their cell cycle, and upon division followed the daughter cells over a number of generations until steady-state is reached. During each cell generation *i*, we evolved the following seven coupled differential equations for cell volume *V*_*i*_, division protein abundance *X*_*i*_, surface area *A*_*i*_, nutrient concentration inside the cell [*n*_*i*_], antibiotic concentration inside the cell 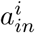, active ribosomes 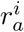, and inactive or antibiotic-bound ribosomes, 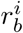.

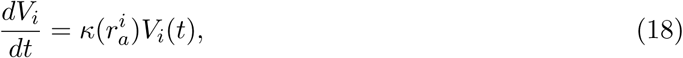

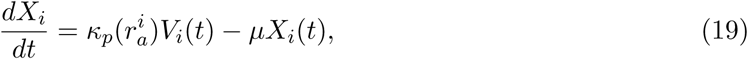

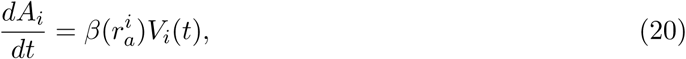

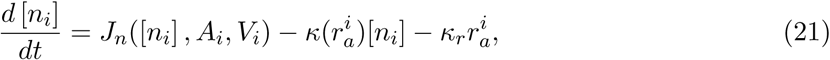

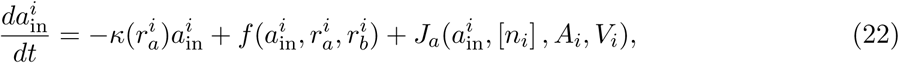

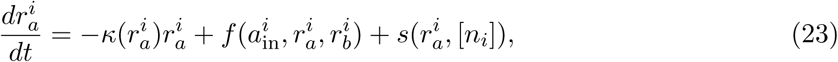

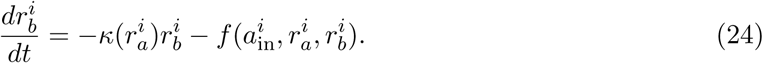

In the above equations, we have (dropping ′*i*′ for simplicity):

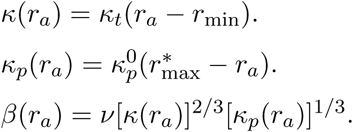

*J*_*n*_(*A, V*, [*n*]) = [*n*_ext_]*P*_in_([*n*])*A/V*, where [*n*_ext_] is the extracellular nutrient concentration, and *P*_in_([*n*]) is the nutrient-dependent inward permeability (Figure 3E)

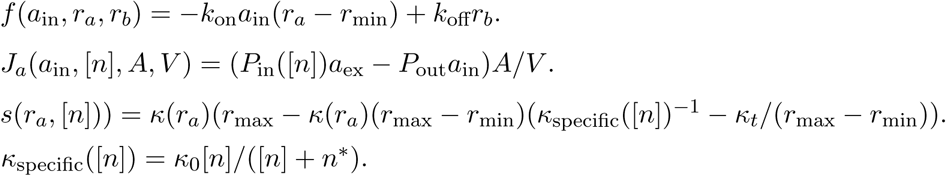

For each cell cycle *i*, Eq. (18)-(24) are evolved for *t* ≤ *τ*_*i*_, where *τ*_*i*_ is the interdivision time for the *i*^th^ generation. Division is triggered when *X*_*i*_ > *X*_0_, with *X*_0_ a constant. Upon division, we set: 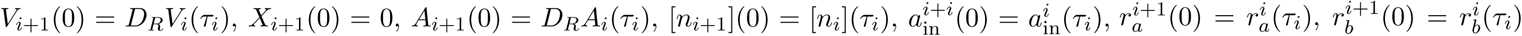, where *D*_*R*_ is a Gaussian random variable with mean 0.5 and standard deviation 0.05. We initialize the nutrient concentration inside the cell ([*n*_*i*_]) close to zero, and calibrate the extracellular nutrient concentration [*n*_ext_] to reach the growth rate of the medium we choose to simulate. Over time [*n*_*i*_] reaches the steady-state value *n*^∗^*κ*_specific_/(*κ*_0_ − *κ*_specific_), such that *κ* = *κ*_specific_. We run simulations for additional 5h after the nutrient concentration reaches steady-state, to record the average values of cell volume, area, and ribosome concentration. Antibiotic perturbation is applied after 10h from the start of the simulations and continued for another 20h, when we compute the average values for the various cellular variables.

### Proteome sector model

The total mass of division proteins, *M*_*X*_, increases at a rate proportional to the amount of actively translating ribosomes, 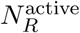,

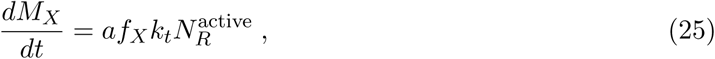

where 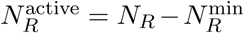, *N*_*R*_ is the total number of ribosomes, 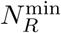 is the number of ribosomes not participating in protein synthesis, *k*_*t*_ is the rate of translation per ribosome, *f*_*X*_ is the fraction of ribosomes devoted to synthesizing *X*, and *a* is the concentration of amino acids. Similarly, 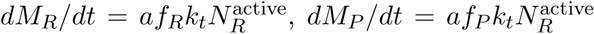, and 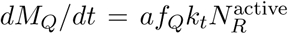, where *M*_*R*_, *M*_*P*_ and *M*_*Q*_ are masses of R, P and Q sector proteins, *f*_*P*_, *f*_*R*_, and *f*_*Q*_ = 1 − *f*_*R*_ − *f*_*X*_ − *f*_*P*_ are the fractions of ribosomes devoted to synthesizing each of these sectors. Therefore, the total dry mass of the cell, *M* = *M*_*P*_ + *M*_*X*_ + *M*_*R*_ + *M*_*Q*_, increases at a rate proportional to the number of active ribosomes,

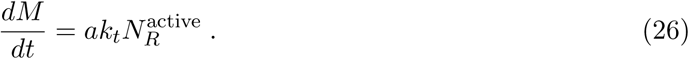

If *m*_*R*_ is the mass of individual ribosomes, we get,

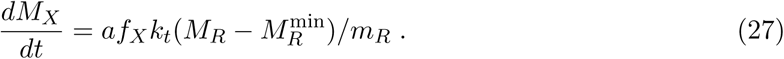

The instantaneous mass fraction of *X, ϕ*_*X*_(*t*) = *M*_*X*_(*t*)*/M* (*t*), then satisfies:

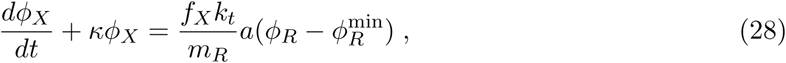

where *ϕ*_*R*_ is the mass of fraction of ribosomes. At steady-state *ϕ*_*X*_ = *f*_*X*_, using the relation: 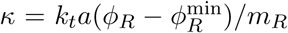. We can rewrite the above equation in terms of the concentration of *X, c*_*X*_ = *ϕ*_*X*_ *ρ*_*c*_*/m*_*x*_, where *ρ*_*c*_ is the mass density of the cell, and *m*_*x*_ is the mass of an individual *X* molecule. This gives us,

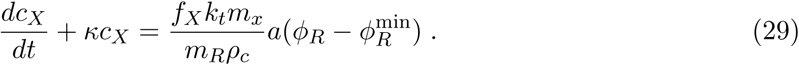

Using *X* = *c*_*X*_*V*, where *X* is total amount of division proteins in the cell, we derive dynamics of division protein accumulation,

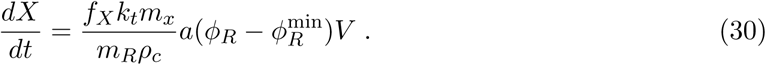

The above equation allows us to identify the division protein production rate as,

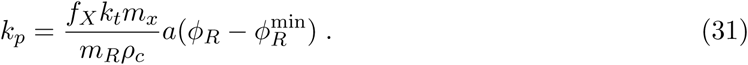

The steady-state concentration of amino acids is determined by the balance between the rate of nutrient influx by transporters and the rate of translation by active ribosomes [26],

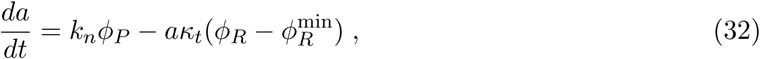

where *ϕ*_*P*_ is the mass fraction of P-sector metabolic proteins, and *κ*_*t*_ = *k*_*t*_*aρ/m*_*R*_. At steady-state, we have 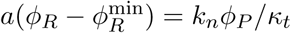. Using 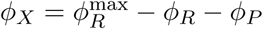, we obtain

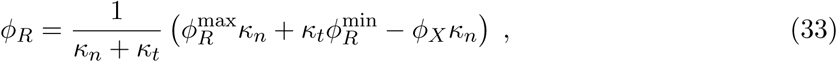

where *κ*_*n*_ = *k*_*n*_*/a*. Therefore, 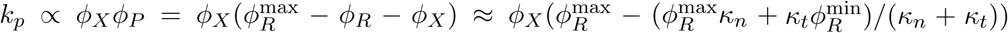, assuming that *ϕ*_*X*_ occupies a small fraction of the proteome. We thus get: 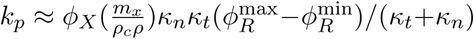. In terms of ribosome mass fraction, the rate of production of division proteins is then given by:

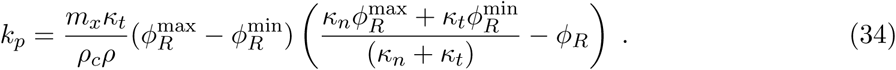

### Parameter determination

We extracted the parameters *κ*_*t*_ and *r*_min_ by fitting the equation *κ* = *κ*_*t*_(*r* − *r*_min_) to the data for growth rate vs RNA/protein ratio [4] (Table 1). Using our theoretical model, we obtained the expression for cell volume *V* as a function of *r* (Eq. (5)), which we fitted to experimental data [4], in order to to extract the parameters 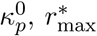 and *μ* (Table 1). For cells under Chloramphenicol stress, the nutrient-dependent parameters *κ*_*n*_ and *δr* were obtained by fitting Eq. (10) and Eq. (11) to the experimental dataset for each nutrient condition (Table 1). From experimental data [4], we estimated the division protein production rate as *κ*_*p*_ = *κ/* ⟨*V*⟩. To determine the permeability of the cell envelope to nutrient and antibiotic transport, we fitted the growth inhibition curves resulting from our simulations to the growth inhibition curves from the data in [4], using a method of least-squares. We find that *P*_in_*/P*_out_ is a function of nutrient quality, and used that as an input to our model simulations. Tables 1 and 2 list a complete set of parameter values used in our model simulations. Note that there are free parameters in the model that are not fixable by fitting experimental data. These include: *X*_0_, *n*^∗^ and *κ*_0_. We do not use *X*_0_ directly, as the value is absorbed in *κ*_*p*_ by renormalizing the number of division proteins by *X*_0_ i.e. *X* (*t* = 0) = 0 and *X* (*t* = *τ*) = *X*_0_ = 1. We determine the nutrient specific growth rate by treating [*n*_ext_]*/n*^∗^ as a fitting parameter. To this end, we arbitrarily pick the values for *n*^∗^ and *κ*_0_ and let the nutrients inside the cell to reach steady-state, *d*[*n*]*/dt* = 0. The steady-state value for [*n*] depends on [*n*_*ext*_] and determines the nutrient-specific growth rate *κ* = *κ*_specific_ = *κ*_0_ [*n*] /(*n*^∗^ + [*n*]). For each growth medium, we tune the value of [*n*_*ext*_]*/n*^∗^ such that it results in the value of *κ* equal to the growth rate reported in experiments.

## ACKNOWLEDGMENTS

We thank Suckjoon Jun lab (UCSD) for providing single cell shape data for *E. coli*, and Guillaume Charras for many useful discussions. SB acknowledges funding from EPSRC grant EP/R029822/1, Royal Society grants URF/R1/180187 and RGF/EA/181044.

## AUTHOR CONTRIBUTIONS

SB and DS conceived and designed research. DS, NO and SB developed theory. DS performed simulations and analysed data. DS and SB wrote the paper.

---

**Figure S 1.**
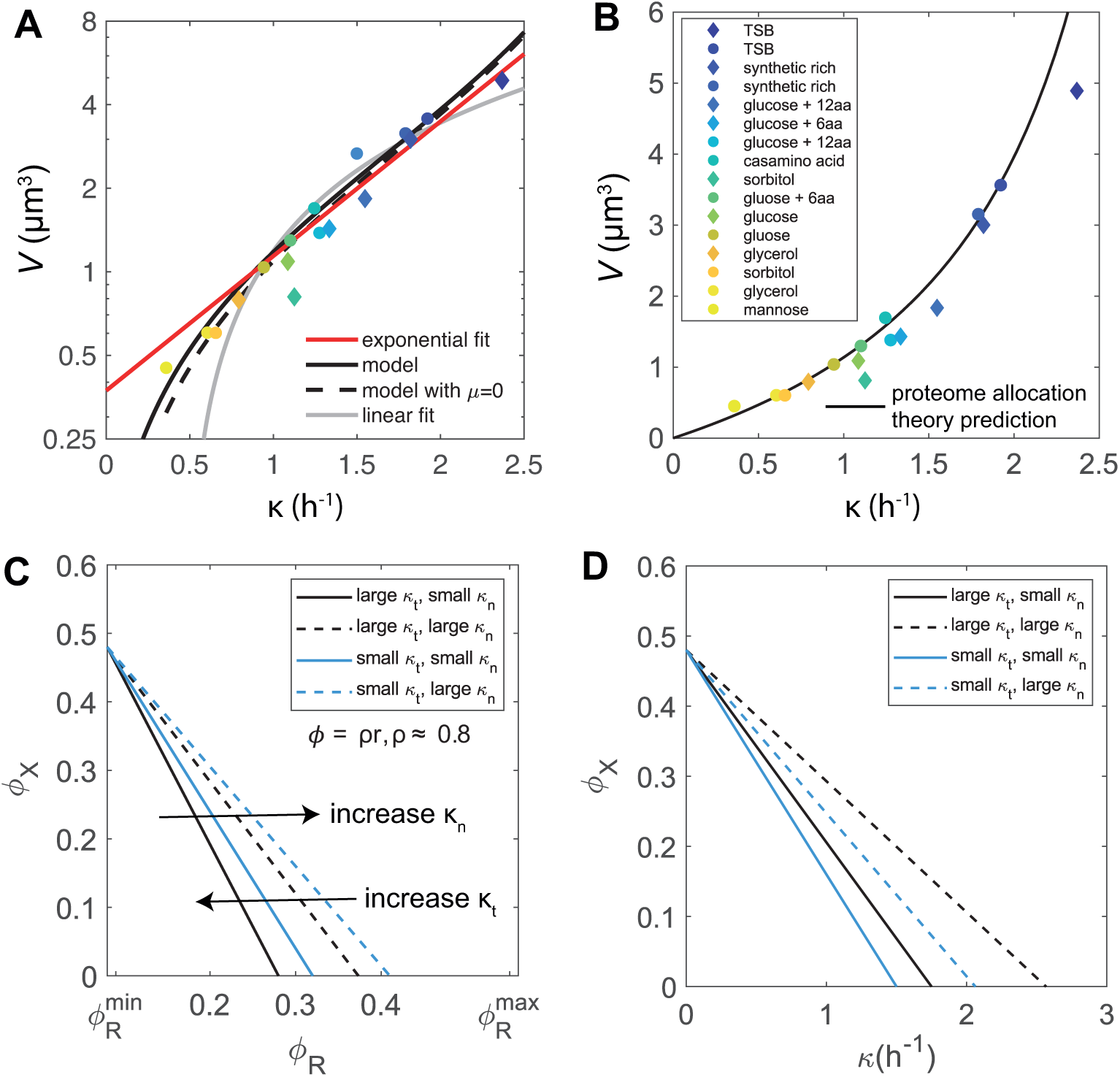
Predictions of the proteome allocation theory. **A**. Cell size vs growth rate, plotted in semi-log scale. **B**. Theoretical prediction for cell size using: 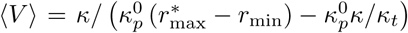, where *r*_max_ depends on the nutrient specific growth rate: 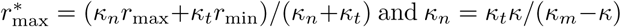 and *κ*_*n*_ = *κ*_*t*_*κ/*(*κ*_*m*_−*κ*). **C**. Dependence of the division protein mass fraction *ϕ*_*X*_ on the ribosome mass fraction *ϕ*_*R*_. Tradeoff between *ϕ*_*X*_ and *ϕ*_*R*_ can is modulated by translational (*κ*_*p*_) and nutrient (*κ*_*n*_) perturbations. *ρ* is a conversion factor between ribosome mass fraction and RNA/protein ratio, *ρ* ≈ 0.8 [36], 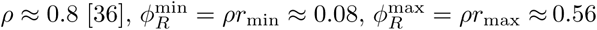. **D**. The same perturbations as in **C**, showing the relationship between *ϕ*_*X*_ and growth rate.

**Figure S 2.**
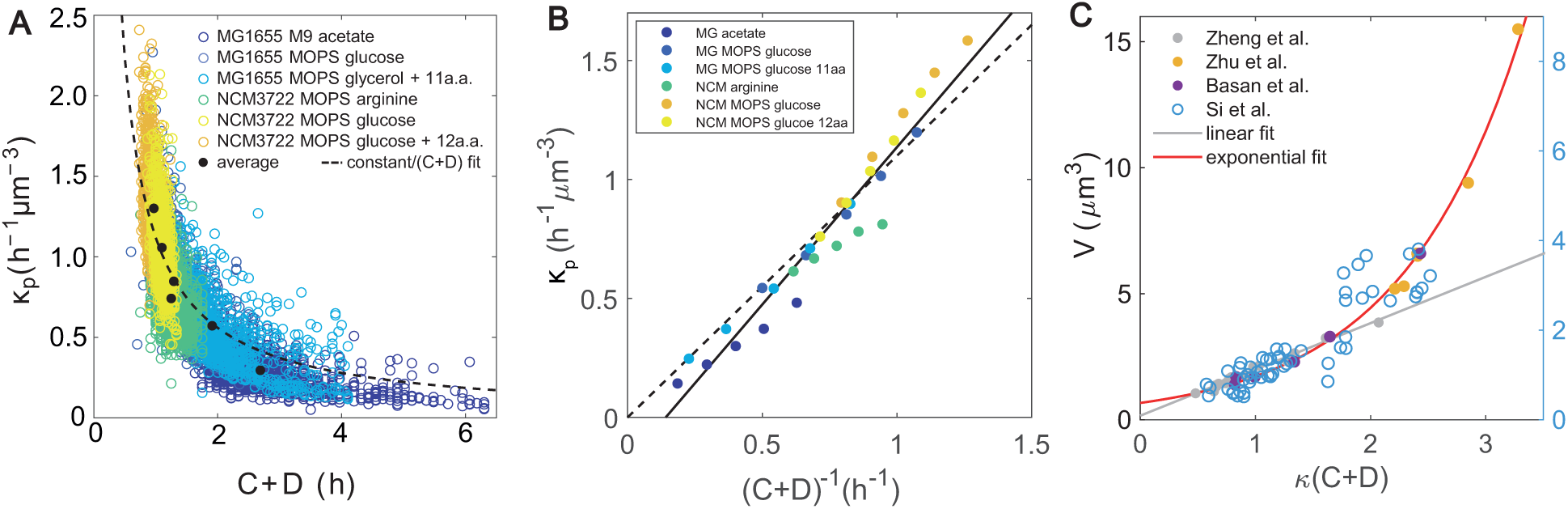
Relationship between cell size, division protein production rate and C+D period. Related to Figure 1. **A**. Scatter plot showing the negative correlation between *κ*_*p*_ and *C* + *D*, with data taken from Ref. [20]. *κ*_*p*_ is estimated using the relation *κ*_*p*_ ≈ *κ/V*. Dashed line is the best fit of the predicted relation *κ*_*p*_ ∝ (*C* + *D*)^−1^, suggesting that measurements of *C* + *D* period could reliably predict the production rate of the division proteins. **B**. Estimating *κ*_*p*_ from measurements of the *C* + *D* period. Dashed line is a fit through the origin as predicted by [48] and solid line is unconstrained linear fit. For each growth condition we removed the outliers that are more than three standard deviations away from the mean and then binned the data. **C**. Relationship between cell size and *κ*(*C* + *D*). The gray line is a linear fit to data from Zheng et al. [48]. The red line is an exponential fit to data from Zheng et al. [48], Zhu et al. [62] and Basan et al. [7]. Data from Si et al. [4] is shown on y-axis to the right. We observe that a linear fit works well for *κ*(*C* + *D*) < 2.

